# Context-dependent effects of microbial inoculation on sagebrush seedlings during drought stress

**DOI:** 10.1101/2025.01.30.635530

**Authors:** Jacob A. Heil, Mathew Geisler, Maddy Skinner, Chadwick DeFehr, Adedotun Arogundade, Bruce Finney, Leonora S. Bittleston

## Abstract

The leaf microbiome interacts with its plant host and can have critical positive and negative effects on plant health. However, the interactions between leaf microbiomes and host plants remain understudied, particularly in reference to context-dependency, e.g., during drought conditions. Plants naturally exist in environments with ever-changing weather and epiphytic foliar microbes are often exposed to extreme conditions. Little is known about how different environmental contexts can alter the effects of phyllosphere microbes on their plant hosts. In this study, we measured the response of seedlings to microbial inoculations and drought conditions. We used *Artemisia tridentata* subsp. *tridentata* (hereafter, sagebrush), a foundation species in the vast and critically threatened sagebrush steppe ecosystem. We grew sagebrush plants in growth chambers from sterile seed and after 6 months the seedlings were inoculated in one of four inoculation treatments: (1) no inoculant, (2) sterile water, (3) the microbes washed from the surface of the leaves (a whole natural community), and (4) the single species, *Bacillus amyloliquefaciens*. We then exposed the seedlings to two environmental treatments: regular watering and a tapered drought. We found context-dependent effects of inoculation on seedling health for both of our inoculation microbial treatments. For example, both inoculants had a negative effect on host plant photosynthesis, which was lessened drought conditions. Furthermore, seedlings inoculated with *B. amyloliquefaciens* had increased leaf nitrogen levels, and the highest survival under drought conditions. Our study shows that seedling health and survival can be influenced by leaf-associated microbes, and is dependent on environmental context.

## Introduction

The microorganisms living both on and in plant leaves (phyllosphere) influence plant health, with effects ranging from pathogenic to mutualistic (e.g., growth promotion, nutrient acquisition, disease resistance) (Stone, Weingarten, and Jackson 2018; Leveau 2019). Plant-microbiome interactions are often context-dependent: they can switch from positive to negative depending on environmental conditions, e.g., *Epichloë* spp. interactions with plants range from positive to negative based on nutrient availability (Stengel et al. 2022). The role of environmental context is critical in understanding plant-microbiome interactions, and it is likely to be exacerbated by rapidly changing climatic conditions (Trivedi et al. 2022). To fully characterize the interactions between a plant and its microbiome it is necessary to account for context, so greenhouse or growth chamber studies where individual plants are inoculated with microbes should include multiple inoculant groups that are each subjected to various environmental contexts to determine if inoculants interact differently with plants in different environments.

Ongoing climate and environmental changes are modifying the contexts in which many plants and microbes have historically interacted. Many ecosystems are increasingly being affected by the environmental stress brought on by drought. Drought modifies the climatic conditions in which microbes live as well as the physiology of plant hosts. Common responses of plants to drought include the closure of stomata, inhibited foliar and stem growth, and changes in root growth strategy (Basu et al. 2016). Closure of stomata leads to increased water retention, decreased photosynthetic activity, and decreased nutrient uptake (Gupta, Rico-Medina, and Caño-Delgado 2020). Microbe-plant associations, such as root-associated arbuscular mycorrhizal fungi (AMF), can promote plant health during drought stress, including in the sagebrush system (Augé, Toler, and Saxton 2015; Geisler, Buerki, and Serpe 2023). Little is known about how the leaf microbiome affects plant physiology during drought stress, but various mechanisms have been proposed as potential functions contributing to plant success during drought (Stone, Weingarten, and Jackson 2018). *Pseudomonas* species, commonly found on leaves, readily form biofilms which may hold moisture in or close to the leaf (Stone, Weingarten, and Jackson 2018). UV resistance may also be a function of biofilms or of pigmented yeasts such as *Aureobasidium pullulans* (Stone, Weingarten, and Jackson 2018; Campana, Fanelli, and Sisti 2022). Plant growth promoting bacteria, including *Bacillus amyloliquefaciens*, are commonly found on leaves and may support plant growth during drought stress through production of the volatile organic compounds acetoin and 2,3-butanediol that induce stomatal closure (Stone, Weingarten, and Jackson 2018; Wu et al. 2019; Ryu et al. 2003). In terms of nutrient acquisition, bacterial nitrogen fixation from phyllospheric bacteria has been observed in Holm Oak trees during drought stress (Rico et al. 2014). Other proposed functions of phyllospheric microbes include heat dissipation and production of abscisic acid to promote stomatal closure (Stone, Weingarten, and Jackson 2018).

For plants growing in arid environments, it is important to consider the response of microbial physiology to drought conditions. Bacterial drought tolerance is facilitated by the formation of endospores as a store of nutrients and the production of osmoprotectant proteins (Sunde et al. 2009; Lebre, De Maayer, and Cowan 2017). Fungi, such as *Aureobasidium pullulans*, regulate internal cation concentrations to survive fluxes in salinity that can result from changes in precipitation (Kogej et al. 2005). Phyllosphere community structure is also affected to some degree at both the levels of host plant species and plant individual (Sanders-Smith et al. 2020; Singh et al. 2023). Persistence of microbial species on specific host plant or in a specific population of plants is conditioned by both plant physiology and the weather (Debray et al. 2022; Heil et al. 2024). For epiphytic microbes, resource availability on the leaf surface is especially important for survival; a factor which is mediated by leaf surface structures (Leveau 2019; Yan et al. 2022). For example, leaf surface heterogeneity accommodates micro-scale pooling of water, creating reservoirs of water and nutrients necessary for bacterial survival (Grinberg et al. 2019). Phyllosphere community structure is influenced by precipitation levels and individual microbial species have differential responses to drought conditions (Bechtold et al. 2021; Heil et al. 2024). Because the survival and proliferation of microbes in the phyllosphere is context dependent, it can be difficult to understand the effects that phyllosphere microbiomes have on their host plant during drought.

*Artemisia tridentata* subsp. *tridentata* (hereafter, sagebrush) is a foundation species in the threatened sagebrush steppe ecosystem that spans most of the western United States (Crist et al. 2019). As drought persists for much of the year in many sagebrush locales, drought stress may even be a normal physiological state or at least a normal seasonal state for many sagebrush populations. Sagebrush have demonstrated the ability to regulate cellular water potential, maintain turgor pressure as well as photosynthesis, and continue root growth during drought (Reed and Loik 2016; Evans et al. 1992; Bassirirad and Caldwell 1992). However, decreased nitrogen uptake, osmotic potential, and leaf solutes have also been recorded (Evans et al. 1992; Bassirirad and Caldwell 1992). Additionally, the degree of drought resistance may be dependent on environmental context and the subspecies (Kolb and Sperry 1999). Recently the fungal phyllosphere community was described for sagebrush throughout the course of a year (Heil et al. 2024). Despite this in-depth characterization over time, we do not know how phyllosphere communities may affect the sagebrush host and if their effects are dependent on environmental context. To address this knowledge gap, we conducted a growth chamber experiment with sagebrush grown from seed and subjected to different environmental conditions and types of microbial inoculants.

Foliar inoculation studies are sparse, especially with regard to naturally-sourced whole communities (Busby et al. 2022). The effects of the microbiome on plant health in these studies range from positive effects, such as pathogen protection and increased growth, to detrimental effects, such as decreased germination and decreased growth. In one rare study showing positive effects, a functionally extinct native Hawaiian mint was protected from powdery mildew by transplanting the microbiome of another mint species (Egan et al. 2021). There may be trade-offs in using whole communities vs. single species inoculants for foliar inoculation. While it is easier to identify the function of a single species as compared to a whole community and thus avoid pathogens, the behavior of single species in monoculture may be different from when it is part of a whole community (Herath 2013). Communities of microbes have been shown to promote plant growth and mitigate disease to a greater extent than monocultures of beneficial bacteria (Herath 2013; Chock, Hoyt, and Amend 2021). This is not to say that single-species inoculants cannot be beneficial. The leaf associate *Bacillus amyloliquefaciens* has been extensively studied as a root inoculum conveying drought resistance, halotolerance, nutrient acquisition, disease mitigation, and more (Sheteiwy et al. 2021; Bisht, Mishra, and Chauhan 2020; Luo et al. 2022). As a foliar inoculant, *B. amyloliquefaciens* has also been shown to mitigate the effects of fungal pathogens (Chien and Huang 2020; Lee and Ryu 2016; Abdelkhalek et al. 2022).

Drought is a common condition for many plant species, including the sagebrush living in much of Western North America. In this study, we inoculated sagebrush leaves with either a single-species or whole-community microbial inoculum to determine the effect on plant health with and without drought conditions. We grew sagebrush in growth chambers from seed sourced from a site in the foothills outside of Boise, ID. Our inoculations included a whole natural microbiome collected directly from wild plants and a single-species inoculant of *B. amyloliquefaciens,* both sourced from the same site as the seeds. We aimed to determine if a natural microbiome conferred health benefits to sagebrush with or without drought conditions. We hypothesized that if either inoculum was helpful to plants under either set of environmental conditions we would observe positive plant health responses such as increased survival, higher photosynthetic activity, growth of above and below ground plant tissue, and increased nitrogen percentage in the leaves. Conversely, if either inoculum was pathogenic we would see a negative effect on our response variables. If the interactions are context dependent, then we would expect to see different responses depending on the environmental conditions. The results of this study inform our understanding of phyllosphere function and the context-dependency of microbiome effects on plant health.

## Materials and Methods

### Experimental design

For this experiment we grew 80 sagebrush test plants from seed collected from a single mother plant living in a sagebrush stand adjacent to the BSU Lower Weather Station (43.6885278, -116.16991; McNamara 2018). To grow the seedlings we filled 164 mL SC10 conetainers with a soil mixture (one part conditioners (1:2:1 volcanic cinder:vermiculite:peat moss) and one part mix (1:1 top soil:compost)) developed for sagebrush by Martinez et al. (Martinez et al. 2023). We designated 10 test plants to each of 8 treatment groups, where they were labeled using unique identifiers. For the duration of the experiment, we grew the test plants in growth chambers (Hoffman Mfg. Inc., Corvallis, OR, USA; SG2-22)at a constant temperature of 25°C with 18 hours of light and 6 hours of dark per 24-hour period. Inside the growth chamber, we kept the plants in a 4×5 grid for each treatment group and rotated the plants on a weekly basis to remove potential positioning bias. The experiment lasted for six weeks from May 25 – June 30 2023, with no photosynthesis measurements taken at the final date due to technical malfunction.

For our experimental design we employed a full factorial design with eight treatment groups: watered-non-inoculated (WN), watered-sterile (WS), watered-whole-community (WW), watered-bacteria (WB), drought-non-inoculated (DN) drought-sterile (DS), drought-whole-community (DW), and drought-bacteria (DB) (Figure 1a). In this design we had two watering regimes, watered (W) and drought (D). To simulate the amount of rainfall that plants would receive in a natural site with an intermediate amount of precipitation (Engel et al. 2024) we watered the plants in the watered (W) groups twice a week with 9 mL of sterile de-ionized (DI) water and the plants in the drought groups with a tapered watering schedule starting at 9 mL of sterile DI water twice in the first week, with each following week decreasing by 2 mL until reaching no watering to simulate drought conditions. For inoculation we had four different inoculant types. We did not inoculate plants in the non-inoculated groups (N). For the sterile water groups (S) we inoculated all plants with autoclaved DI water, for the whole natural community groups (W) we inoculated each plant with a water suspension of microbes washed directly from about 2 g of freshly collected sagebrush leaves, and for the *B. amyloliquefaciens* group (B) we inoculated all plants with a live suspension of ∼250,000 cells/mL of *B. amyloliquefaciens* in water.

**Figure 1.**
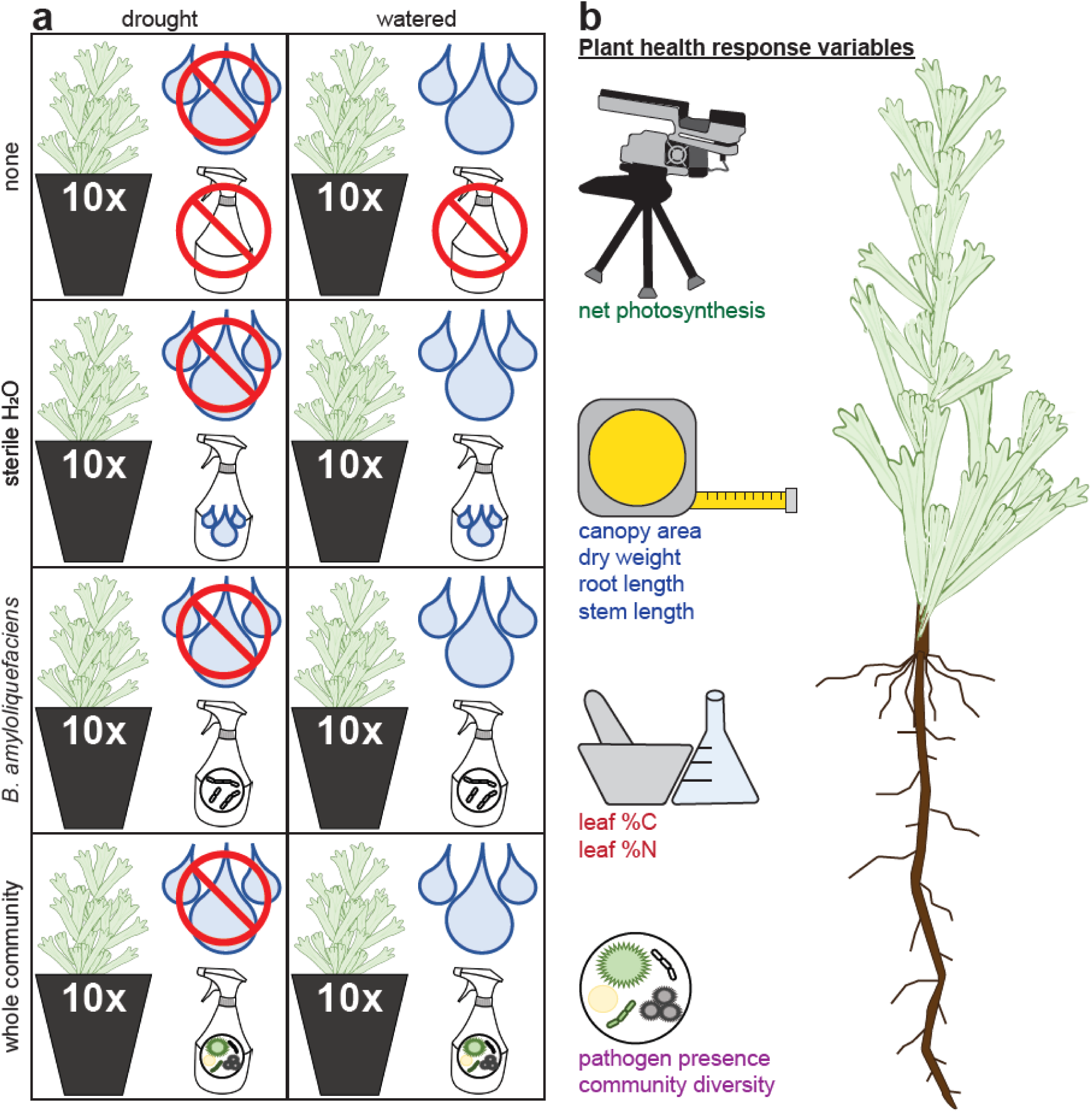
Experimental design. a. We split 80 plants evenly between eight treatment groups using four different inoculants (none, sterile water, *B. amyloliquefaciens*, whole natural community) and two different environments (drought, watered). b. We quantified plant health by collecting 9 response variables.

### Inoculation

To inoculate plants we used sterilized plastic tents as inoculation chambers. For sterilization, we sprayed the tents with 10% bleach, wiped them down, allowed them to sit for 10 minutes, and then wiped down the tents with 70% ethanol followed by a UV light application for 10 minutes. Each inoculant was prepared the day of inoculation. We sterilized the spray bottles by applying 10% bleach for 10 minutes, rinsed them twice with sterile water, then rinsed them with 70% ethanol for another 10 minutes, and then did two final rinses with sterile water. We sprayed every bottle at each sterilization step to ensure the spray hose and nozzle was thoroughly sterilized. For the single species inoculant we chose a bacterial endophyte previously recovered from sagebrush leaves, *Bacillus amyloliquefaciens* (GenBank #PQ660773). We prepared the single species inoculant by growing *B. amyloliquefaciens* in potato dextrose broth on a shaker at 200 rpm at room temperature for 2 days, then transferred the inoculated broth to a falcon tube to be centrifuged at 10,000 rpm for 5 minutes. We removed the supernatant and resuspended the bacterial pellet in 45 mL of sterile DI water, vortexed for 2 minutes, and centrifuged again. We then removed the supernatant and resuspended the pellet in 150 mL of sterile DI water before staining with cotton blue dye to count the number of cells in a hemocytometer. We used the average cell count from the outer four squares of the hemocytometer to dilute the concentration of the inoculant to 500,000 cells/mL. To develop the whole community inoculant we determined the volume of leaves that would reliably inoculate Potato Dextrose Agar (PDA) plates with a natural community. We focused on a volume of leaves to replicate a natural community in a given area of leaf surface. We gathered approximately 25 mL of sagebrush leaves from the same stand where we collected the seeds for our test plants into 3 separate falcon tubes and filled the tubes with 25 mL of sterile DI water before vortexing for 2 minutes. We filtered the water and sagebrush leaf solution into a new tube, and the leaves were then washed with the same process until 150 mL of water was collected. We loaded both inoculant solutions into their designated spray bottles and sprayed onto a PDA plate to test for efficacy at each inoculation date. The sterile water inoculant was simply sterile water in a sterile spray bottle. To inoculate individual plants, we sprayed each plant one time in a sweeping motion from the top of the plant to the bottom, about 6 inches away from the plant, and each spray contained ∼690 μL of inoculant.

For sagebrush leaf sampling, we used sterile technique: sterilizing tents before plants were placed inside to prevent outside contamination, and using clean, gloved hands to handle the plants inside the tents to avoid direct contact. We collected leaves using tweezers that were sterilized with 70% ethanol between each plant and we placed sampled leaves into sterile 1.5mL vortex tubes containing 200 uL of sterile water. To acquire epiphytic microbes, we then vortexed the tubes for 2 minutes before removing the water and dispensing it onto a PDA plate and spreading the water across the plate using 3-5 sterile glass beads then growing the culture plates at room temperature for one week. To acquire endophytic microbes, we removed the leftover leaves in the tubes, cut them into 1 mm x 1 mm pieces, and loaded them into tea strainers for surface sterilization. To sterilize the leaf pieces, the leaves were first washed in 8.5% tween for 2 minutes, then washed for 2 minutes with 70% ethanol, then twice for 2 minutes in sterile water. After the four washes were complete, we removed the leaf pieces from the tea strainers and placed them evenly apart on PDA plates to allow any endogenous microbes to grow at room temperature for one week.

### Sampling procedures

To determine the response of test plants to inoculation and environmental conditions we measured variables related to four plant health categories: survival, photosynthetic activity, growth, and nutrient content. Additionally, we described the bacterial and fungal communities. To quantify photosynthetic activity we measured net photosynthesis at each of five sampling dates using a LI-6400-40 leaf chamber fluorometer connected to a LI-COR LI-6400XT portable photosynthesis system (LI-COR Inc., Lincoln, NE, USA) and following the procedure for sampling sagebrush outlined in Geisler, Buerki, and Serpe 2023 and including a sterilization step in between plant where we sterilized the measurement chamber with 70% ethanol. We derived a mortality index from photosynthetic activity by classifying a plant as dead when the plant’s net photosynthesis fell below 2µmol^-2^s^-1^ and was never measured above 2 again in subsequent samples. Additionally, we quantified the %C and %N in the leaves at the final sample date at the Idaho State University Stable Isotope Laboratory using a Costech ECS 4010 elemental analyzer interfaced with a Thermo Delta V Advantage continuous flow isotope ratio mass spectrometer (Thermo Fisher Scientific, Waltham, MA, USA) following the procedure outlined in (Heil et al. 2024). To quantify plant growth we measured the change in canopy area throughout the experiment, the final weight of the plants, and the length of the root and the shoot of the plants at the end of the experiment. For canopy area and the lengths of roots and shoots we took pictures of each next to a ruler and used ImageJ (Schneider, Rasband, and Eliceiri 2012) to calibrate the scale to 1 cm on the ruler and then measure the plant parts by manually drawing the objects to be measured over them. We determined the final weight of plants by drying each plant at 60°C for 48 hours and then weighing on a calibrated scale.

To characterize the microbial communities we extracted DNA from samples of three leaves from each test plant at the final sample date using the ZymoBIOMICS 96 MagBead DNA Kit (Zymo Research Corp, Irvine, CA, USA; Cat # D3408) and determined DNA quality using the AccuClear Ultra High Sensitivity dsDNA Kit (Cat # 31028). We assessed the efficacy of sequencing by using a negative control (DNAse free water) and for the positive control we used ZymoBIOMICS Microbial Community Standard (Cat # D6300). Amplicon library preparation and sequencing were performed at the Environmental Sample Preparation and Sequencing Facility (ESPSF) at Argonne National Laboratory for amplicon sequencing on a MiSeq. We targeted bacteria using chloroplast-excluding primers (799F and 1115R) for the V5-V6 region of the 16S rRNA gene (Redford et al. 2010; Laforest-Lapointe, Messier, and Kembel 2017), and we targeted fungi using the ITS1F and ITS2 primers (Manter and Vivanco 2007). We used QIIME2 version 2022.2 (Bolyen et al. 2019) and the DADA2 plugin (Callahan, McMurdie, and Rosen 2016) to check the quality of our sequences, remove chimeras, and generate amplicon sequence variants (ASVs). Our 80 samples contained 1,950 16S ASVs and 1,075 ITS ASVs. We assigned taxonomy following the method described in (Heil et al. 2024), where we first used rBLAST to match ASVs with sequences from our in-house sagebrush phyllosphere database, which we curated from ITS and 16S DNA sequences extracted from fungal and bacterial cultures of sagebrush leaf associates, and then against the NCBI’s ITS RefSeq Fungi or core nucleotide database for 16S (O’Leary et al. 2016).

### Data analysis

All downstream data analysis was done in the R program version 4.4.0 (R Team 2023). To model the response of various plant health factors to test for effect sizes of predictor variables we used Bayesian generalized linear mixed models (glmms) using the *brms* package and weakly informative priors (Bürkner 2017). Each model contained the categorical predictor variables of environment (watered or drought) and inoculant (none, sterile water, *B. amyloliquefaciens*, and a natural whole community) as well as an interaction term to determined interactive effects of both predictors. For the net photosynthesis model we used the Gaussian distribution and the random intercepts of plant individual and sampling date to account for any potential error introduced by the LICOR LI-6400XT. For the canopy area model we used the Gaussian distribution and the random intercept of plant individual. For the end-point response variables %C and %N we used a beta distribution and for all other end-point response variables we used Gaussian distributions. We constructed all glmms with the *brms* package (Bürkner 2017), this package uses a Markov Chain Monte Carlo (MCMC) sampler. All of our models employed weakly informative priors, a burn-in period, and chain specifications. To judge model fit we checked effective sample size and model convergence, indicated by Gelman–Rubin statistics close to 1 and stable, well-mixed chains (Gelman and Rubin 1992).

For both the bacterial and fungal microbial communities we used the *vegan* package (Oksanen et al. 2022) to calculate Bray-Curtis dissimilarity using the *vegdist* function, dbRDA using the *dbrda* function, and the *anova.cca* function to assess statistical significance of the terms from the dbRDA model. For dbRDA we used environment and inoculant as predictor variables and plant individual as the grouping variable. We determined differential abundance of genera using the *ANCOMBC* (Lin and Pedadda 2020) package. Our ANCOM-BC model included the relative abundance of taxa grouped at the genus level and environment and inoculant as predictor variables.

## Results

### Microbial community

Leaf microbial community composition was shaped by the inoculant type for both the bacterial and fungal communities (Figure 2a,d), despite the presence of non-target microbes. We began with sterile conditions, using sterilized seeds and soil, and, while we maintained as sterile conditions as possible, we observed non-target bacteria and fungi which were not part of our inoculations and were present in all treatment groups (Figure 2b,e). It is very difficult to maintain sterile growth conditions in growth chambers without air filters that need to be opened and closed for watering the plants, and microbes from the surrounding environment colonized our seedlings. However, the presence of these additional bacteria and fungi made our groups more similar to each other, so the differences we found in response to the environmental treatments and inoculant types should be considered conservative estimates. From our DNA results we determined that the fungal genera *Cladosporium*, *Penicillium*, and *Acremonium*, which are molds commonly found in indoor environments, were present in all samples (Figure 2e). Corroborating the DNA results, we also noted a few morphologically distinct fungi present at all time points in our leaf cultures (Figure S1).

**Figure 2.**
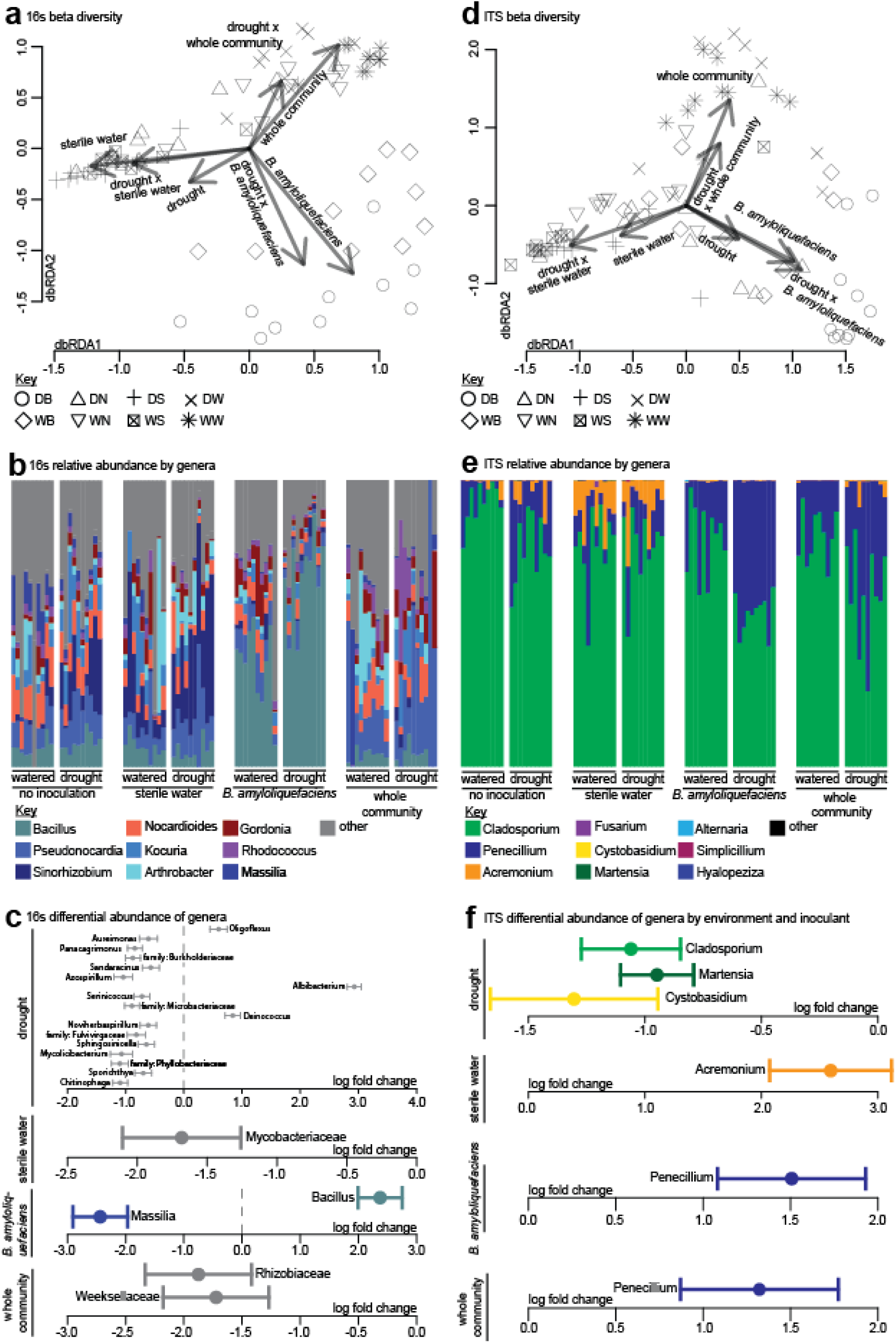
Inoculant type affects the bacteria (16S) and fungi (ITS) of the sagebrush seedling leaf microbiome. a) Distance-based redundancy analysis (dbRDA) using 16S Bray Curtis dissimilarity, with arrows for significant predictor variables overlaid. Point shape indicates treatment group. b) Relative abundance of the top 9 bacterial genera across treatment groups. c) Log fold change of bacterial taxa with significant differential abundances in treatments as determined by ANCOM-BC. d) dbRDA using ITS Bray Curtis dissimilarity, with arrows for significant predictor variables overlaid. Point shape indicates treatment group. e) Relative abundance of the top 9 fungal genera. f) Log fold change of fungal taxa with significant differential abundances in treatments as determined by ANCOM-BC.

To determine the effects of inoculant type and environmental conditions on the microbial communities of our sagebrush seedlings, we measured beta diversity for both the bacterial and fungal communities as well as differential abundance among all genera across treatments. For 16S beta diversity, we found that community similarity was influenced by inoculant type (F = 11.489, p = 0.001, Figure 2a), watering regime (F = 5.750, p = 0.001), and the interactive effect of inoculant and watering (F = 2.169, p = 0.005). In the 16S community, the genus *Bacillus* had the highest relative abundance (Figure 2b) and was differentially abundant in plants inoculated with *B. amyloliquefaciens* (lfc = 2.37, p = 3.07e-08, Figure 2c), confirming that our inoculations led to successful colonization of leaves. We found an overall decrease in differentially abundant genera during drought conditions, with 14 genera having lower relative abundances versus only three with higher relative abundances in drought treatment plants (Figure 2c). In terms of our inoculations, genera from the family *Mycobacteriaceae* had significantly lower differential abundance in plants inoculated with sterile water (lfc = -1.68, p = 1.21e-03), the genus *Massilia* had significantly lower differential abundance in plants inoculated with *B. amyloliquefaciens* (lfc = -2.43, p = 2.86e-06), and the genera *Sinorhizobium* (lfc = -1.87, p = 1.62e-04) and *Empedobacter* (lfc = -1.72, p = 3.62e-04) had significantly lower differential abundances in plants inoculated with a natural whole community.

For ITS beta diversity, we found that community similarity was influenced by inoculant type (F = 2.991, p = 0.001, Figure 2d), watering regime (F = 1.899, p = 0.001), and the interactive effect of inoculant and watering (F = 1.128, p = 0.035). *Cladosporium* and *Penicillium* accounted for over 75% of total fungal abundance in every sample, with *Acremonium* being the third most abundant genus (Figure 2e). For the plants subjected to drought, we found that *Cladosporium* (lfc = -0.97, p =1.85e-05), *Martensia* (lfc = -0.86, p = 5.76e-04) and *Cystobasidium* (lfc = -1.27, p = 7.82e-06) all had lower differential abundances (Figure 2f). We found that *Acremonium* had higher differential abundance in plants inoculated in sterile water (lfc = 2.59, p = 2.59e-05) and that *Penicillium* had higher differential abundance in plants inoculated with both *B. amyloliquefaciens* (lfc = 1.51, p = 6.54e-4) and a natural whole community (lfc = 1.32, p = 4.55e-3). For the *B. amyloliquefaciens*-inoculated plants, the increase in relative abundance of *Penicillium* was paired with a qualitative decrease in the relative abundance of *Cladosporium* and *Acremonium*.

### Photosynthetic activity

We used net photosynthesis to estimate the mortality of plants at each sample date and expressed mortality as a proportion of the total plants in each sample group (Figure 3a). We observed that the watered treatment groups inoculated with sterile water and *B. amyloliquefaciens* had a 90%+ survival rate throughout the experiment. The watered treatment group inoculated with a whole community had 50% mortality at the final time point. Plants from drought treatment groups died faster than most watered plants. Of the drought treatment plants, the *B. amyloliquefaciens*-inoculated plants had the highest survival at 60% while plants inoculated with sterile water or the whole community had 50% survival at the final time point (Figure 3a).

**Figure 3.**
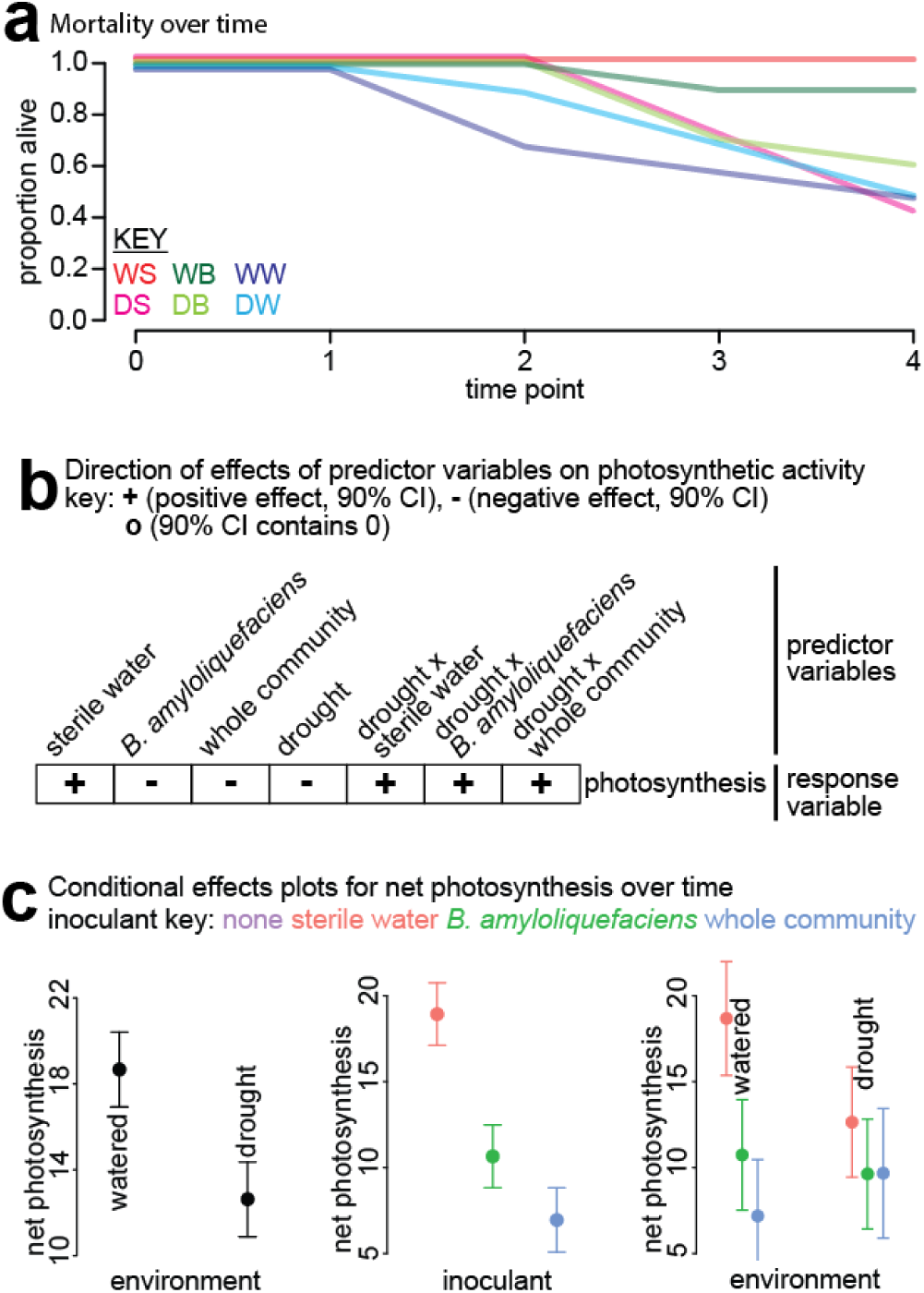
Mortality and photosynthetic activity differ across treatments. Results from measurements of net photosynthesis demonstrating estimated mortality and the effects of inoculants and drought on plant photosynthetic activity. a) Proportion of estimated mortality over time by treatment group (color coded based on the key in the figure: W as a first letter = Watered, D = Drought, S = Sterile water inoculation, B = *B. amyloliquefaciens* inoculation, W as a second letter = Whole community inoculation). b) Direction of effects for predictor variables (columns) for the response variable, net photosynthesis. Plus signs indicate an effect size distribution that is 90% or more above zero. Minus signs indicate an effect size distribution that is 90% or more below zero. c) Conditional effects plots for the photosynthesis model.

Plants inoculated with sterile water maintained higher photosynthetic rates than plants that were inoculated with either *B. amyloliquefaciens* or a whole natural community. After accounting for the random effects of sampling date (sd = 3.68) and the plant individual (sd = 0.70), we found that inoculation with sterile water had a non-zero positive relationship with net photosynthesis (estimate = 7.67, se = 2.00; Figure 3b; Table S1), while inoculation with a whole natural community had a non-zero negative relationship (estimate = -3.81, se = 1.99) and the 90% CI for the effect of *B. amyloliquefaciens* contained zero (estimate = -0.07, se = 2.01). We also found that photosynthesis was higher for plants receiving constant water, while drought had a non-zero negative effect on photosynthesis, as would be expected (estimate = -5.23, se = 2.74). Microbial inoculants tended to decrease photosynthesis under well-watered conditions, but the negative effects of drought and microbial inoculation were lessened by the interaction of the environment and inoculant. The bacterial (estimate = 4.86, se = 2.09) and whole community (estimate = 8.10, se = 2.10) inoculant had higher photosynthesis in drought than expected based on the expected effect of drought (Figure 3b and 3c). We found that the effects of all other predictor variables contained zero within a 90% credible confidence interval (CI).

### Plant growth

We measured plant growth by canopy area (Figure 4a, Table S2) whole plant dry weight (Figure 4a, Table S3) as well as root length (Table S4) and stem length (Table S5). We found that the whole community inoculant strongly reduced canopy area. After accounting for the random effect of plant individual (sd = 3.13), we found that inoculation with a natural whole community had a non-zero negative relationship with canopy area (estimate = -3.37, se = 1.45; Figure 4a,b; Table S2). We measured all other plant growth response variables once at the end of the experiment so we did not account for the random effect of sampling date or individual plant in those models. In contrast to photosynthesis, drought treated plants had greater biomass than regularly watered plants, suggesting that fungal pathogens are consuming plant biomass to a greater degree during regular watering. For whole plant dry weight we found that the drought condition had a non-zero positive relationship with whole plant dry weight (estimate = 0.24, se = 0.10; Figure 3a,d; Table S3); in contrast, inoculants decreased plant weight: both *B. amyloliquefaciens* (estimate = -0.13, se = 0.10) and the natural whole community (estimate = -0.23, se = 0.10) had non-zero negative relationships. Roots were likewise shorter when a natural whole community was inoculated, with a non-zero negative relationship with root length (estimate = -5.23, se = 1.63; Table S4). All other effects of predictor variables in these models contained zero within a 90% CI and so are not discussed further.

**Figure 4.**
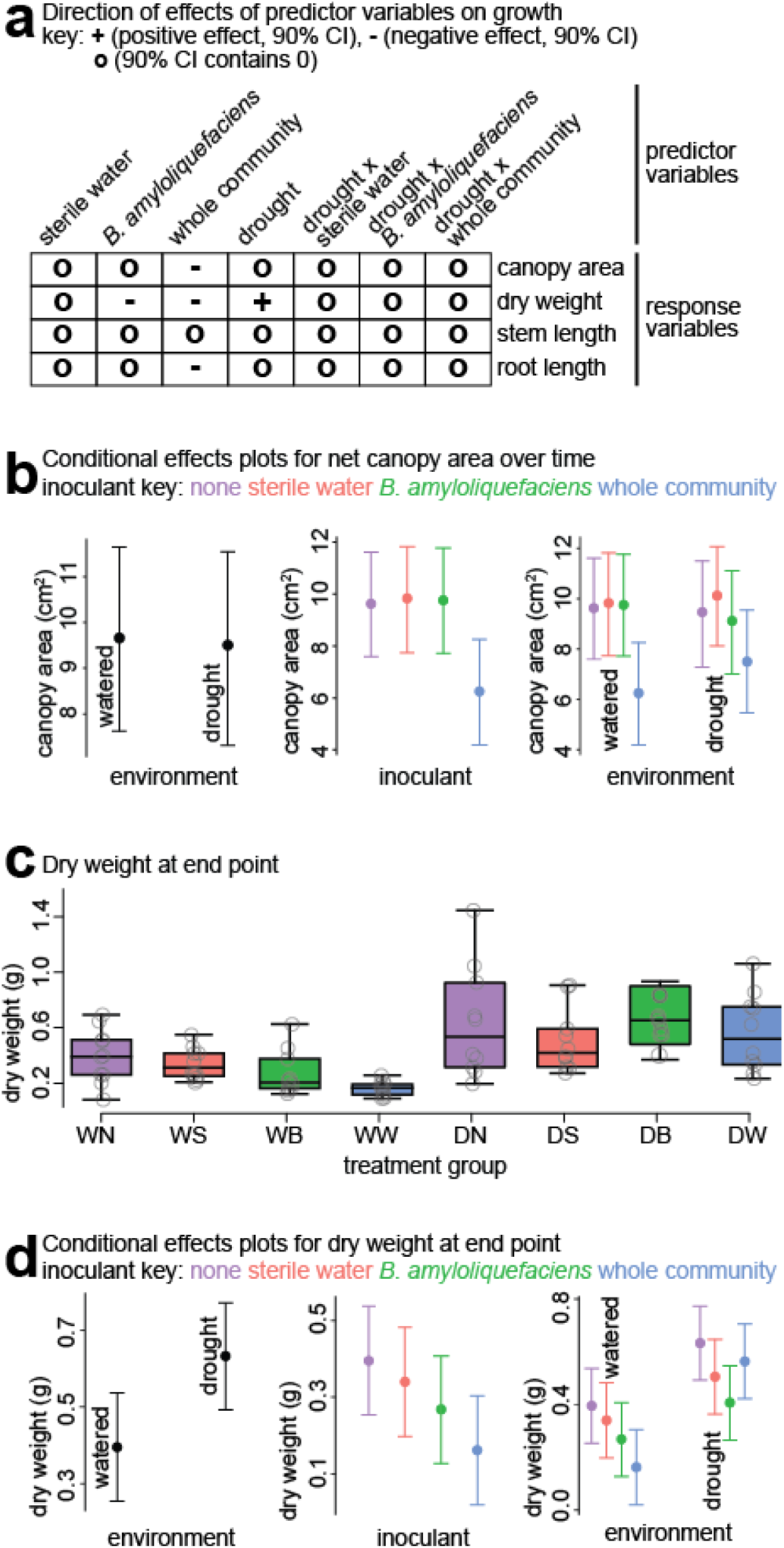
Microbial inoculants had negative effects on plant growth. a) Direction of effects for predictor variables (columns) for two response variables (rows). Plus signs indicate an effect size distribution that is 90% or more above zero, while minus signs indicate an effect size distribution that is 90% or more below zero. Black circles indicate that the 90% CI for the effect size contained zero. b) Conditional effects plots for the canopy area model. c) Raw data for dry weight in each treatment group. d) Conditional effects plots for dry weight model.

### Nutrient content

We measured nutrient content by leaf weight %C and %N (Figure 5a, Table S6, Table S7). We found that the drought condition had a non-zero negative relationship with %C (estimate = -0.09, se = 0.04; Figure 5a,c; Table S6) and that during drought, inoculation with a whole natural community had a positive non-zero relationship (estimate = 0.15, se = 0.06). These results suggest that while drought decreases the amount of carbon in leaves, whole community inoculation can partly offset that effect. For %N, we found that inoculation with *B. amlyloliquefaciens* had a very strong positive non-zero relationship with %N (estimate = 0.15, se = 0.06; Figure 5a,e; Table S7) and inoculation with a natural whole community has a non-zero negative relationship (estimate = -0.09, se = 0.07). The enriching effect of *B. amyloliquefaciens* on N was also reflected in the C:N ratio (mean = 11.46, estimate = -0.14, se = 0.05; Table SX). These results suggest contrasting effects of inoculants: *B. amlyloliquefaciens* increased sagebrush leaf nitrogen percentages, while the whole community decreased them.

**Figure 5.**
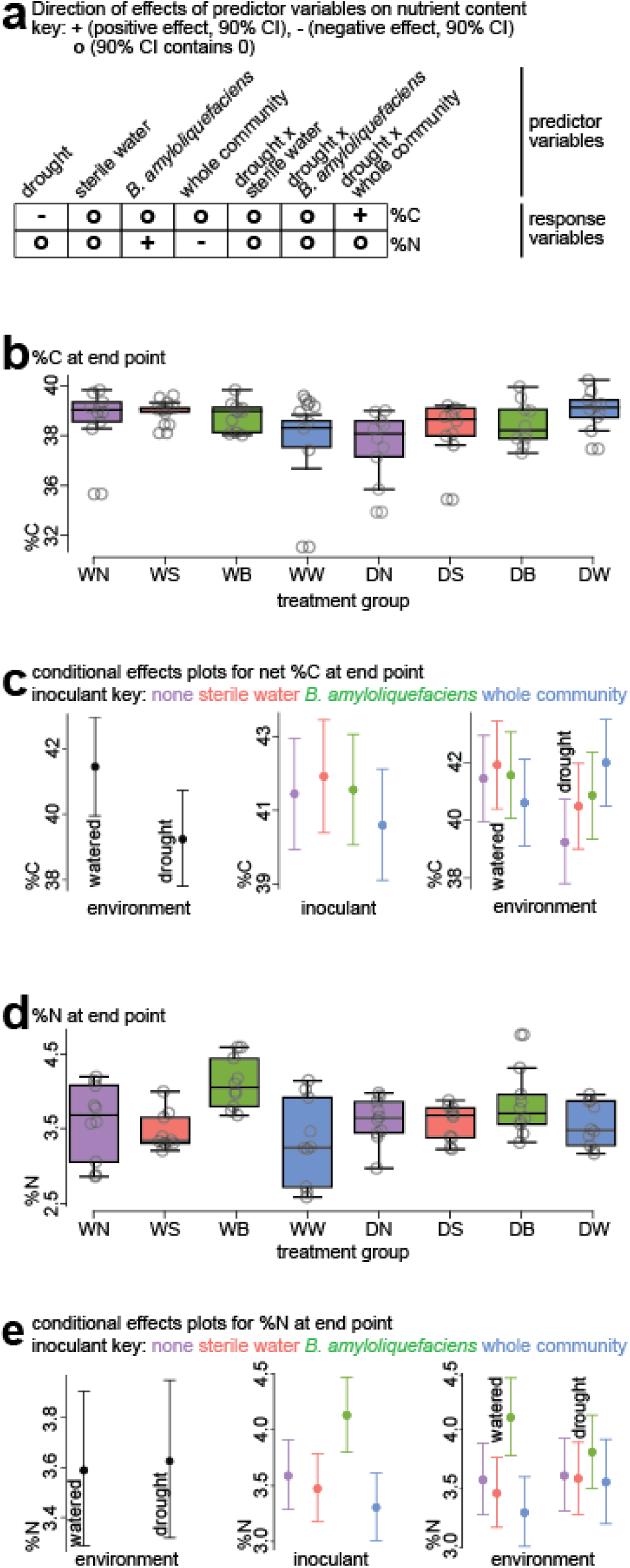
Inoculation with *B. amyloliquefaciens* increased leaf percent nitrogen. Results from models demonstrating the effects of treatment groups on nutrient content. a) Direction of effects for predictor variables (columns) for two response variables (rows). Plus signs indicate an effect size distribution that is 90% or more above zero. Minus signs indicate an effect size distribution that is 90% or more below zero. Black circles indicate that the 90% CI for the effect size contained zero. b) Raw data for %C in each treatment group. c) Conditional effects plots for the %C model. d) Raw data for %N in each treatment group. e) Conditional effects plots for %N model.

## Discussion

In this study we measured both positive and negative plant health responses to microbial inoculants and drought conditions. While inoculations with both the bacterium *B. amyloliquefaciens* and a whole leaf microbial community decreased net photosynthesis, this effect was lessened under drought conditions. Perhaps surprisingly, inoculation with *B. amyloliquefaciens* increased leaf %N by 15% on average. Our analyses were complicated by the presence of additional microbes in all our sagebrush seedlings (Figure 2b,e). Through leaf cultures (Figure S1) and sequencing results we identified the main non-target microbes as fungi from the genera *Cladosporium*, *Penicillium*, and *Acremonium*. Despite this, we noted distinct effects of our inoculants on phyllosphere community structure, mortality, photosynthesis, growth, and nutrient content.

To monitor photosynthesis, we leveraged repeated measurements every two weeks for the duration of the experiment. Plants subject to drought conditions had 50% lower photosynthetic activity than the regularly watered plants. Decreased photosynthesis is a normal response of sagebrush to drought and the ability for the plant to regulate its photosynthetic response is dependent on environmental conditions aa well as the subspecies (Reed and Loik 2016). Plants receiving microbial inoculants likewise had decreased photosynthetic activity. *B. amyloliquefaciens-*inoculated plants had 42% lower net photosynthesis than the sterile-water inoculated plants and whole-community inoculated plants were 62% lower. There were potentially pathogenic bacteria and fungi present on all test plants (Figure 2b and e) which may decrease photosynthesis through destruction of leaf tissue (Najafpour 2012).

Although the leaf communities of our test plants did not resemble natural sagebrush leaf communities (Heil et al. 2024), we observed that community structure was most strongly affected by inoculant but also affected by watering regime and the interaction of inoculant and watering regime. Inoculation with *B. amyloliquefaciens* decreased the relative abundance of bacteria from the genus *Massilia* (Figure 2c). *Massilia* is a common plant root and leaf associate (Ofek, Hadar, and Minz 2012; Xu et al. 2023) and may function as a plant growth promoter and mediator of plant disease (Q. Han et al. 2024; Li et al. 2021). In previous studies, inoculation with *B. amyloliquefaciens* has had both positive and negative effects on *Massilia spp.* abundance (Uwaremwe et al. 2022; Maslennikova et al. 2023; Wan, Zhao, and Wang 2017). The potential pathogen, *Penicillium spp.* was enriched in both inoculant groups. In contrast, other studies have found that *Penicillium spp.* can be suppressed by inoculation with *B. amyloliquefaciens* (Ji et al. 2013; Larbi-Koranteng, Awuah, and Kankam 2020). Little is known about the interactions between individual species in community contexts and our results suggest that they can be variable depending on context. Our inoculants were effective in structuring the leaf communities of our test plants and further research is required to have a full understanding of their specific effects on microbe-microbe interactions and plant health.

Through their influence on community structure, our microbial inoculants may have augmented the pathogenic effect of the leaf microbiome to be greater than in sterile-water inoculated plants. A differential influence on the relative abundance of some fungal species over others may have driven this effect (e.g. both microbial inoculations increased the relative abundance of *Penicillium*, Figure 2f). The decrease in net photosynthesis by both drought and microbial inoculation was dampened when they were combined (Figure 3). Past studies have shown variable outcomes for the interactive effect of plant pathogens and drought on plant health (Milici et al. 2020). Precipitation can cause proliferation and spread of pathogens; however, the detrimental effects of drought on plant physiology may make them more susceptible to the effects of pathogens. The ability of sagebrush to regulate its physiological response to drought (Reed and Loik 2016; Evans et al. 1992; Bassirirad and Caldwell 1992) suggests that the plant’s susceptibility to disease is not strongly increased in drought conditions and that the pathogen-promoting conditions during regular watering cause stronger pathogenicity. From our sequencing data, drought decreased the relative abundance of many bacterial taxa including potential plant pathogens such as bacteria from the family *Microbacteriaceae* (Dworkin et al. 2006) and the genus *Azospirillum* (Tortora, Díaz-Ricci, and Pedraza 2012) (Figure 2c).

In the field, the degree to which sagebrush is affected by drought varies contextually, but sagebrush sometimes demonstrates a capacity to maintain its photosynthetic rates (Reed and Loik 2016). Our results indicate that during drought conditions, the expected effect of drought on photosynthesis was less pronounced in both microbial inoculant groups (Figure 3a,b). *B. amyloliquefaciens* is a common root-associate in plants (Sheteiwy et al. 2021) and the strain used in this study was initially cultured as a sagebrush leaf endophyte. In root inoculation studies, *B. amyloliquefaciens* has demonstrated plant growth promotion and promotion of photosynthetic efficiency by augmenting chlorophyll fluorescence and gas exchange (L. Han et al. 2022; Samaniego-Gámez et al. 2016; Xie et al. 2018). The negative effect *B. amyloliquefaciens* on net photosynthesis was less pronounced than the whole community (Figure 3a), suggesting that there may have been a limited antagonistic effect of *B. amyloliquefaciens* driven by increased relative abundance of some plant pathogens.

For plant growth, we measured a pronounced negative effect of the whole community inoculant on both net canopy area and dry weight as well as a negative effect of *B. amyloliquefaciens* on dry weight (Figure 4). Surprisingly, plants in the drought treatment had increased dry weight. The positive effect on plant weight by drought was more pronounced in the leaves rather than the stem or roots. Every plant had fungi from the genera *Cladosporium*, *Penicillium*, and *Acremonium* (Figure 5b), each of which contain species with the capacity to cause leaf blight and rot (Thomma et al. 2005; J.-X. Han et al. 2023; Rashed 2018). *Cladosporium* as a plant pathogen has been shown to decrease whole plant biomass in timothy grass (Yang and Li 2022). The environmental treatment group that received water may have allowed for higher pathogenicity of one or all of these species and, similarly, the microbial inoculant treatments seemed to act as a catalyst for the foliar pathogens. In studies from other plants, *B. amyloliquefaciens* has shown plant growth promoting properties as well as having antagonistic effects on some fungi (Ngalimat et al. 2021). Therefore, it was surprising to observe that *B. amyloliquefaciens* had a negative effect on plant dry weight, suggesting that it exacerbated the negative effects of the pathogenic fungi. From our community analyses, we could see that *B. amyloliquefaciens* played a role in structuring the fungal community (Figure 2d and e) by increasing the relative abundance of fungi from the genus *Penicillium* in relation to the two other highly-abundant genera *Cladosporium* and *Acremonium*. In this study, opportunistic pathogens likely took hold as the seedlings were growing and long before our treatments began. A promising future direction would be to begin treatment inoculations as the first leaves emerge on the seedlings, to see if early inoculation with more beneficial microbes can inhibit the establishment of ones with negative effects.

We also observed mixed leaf nutrient responses (Figure 5), for example, drought had a negative effect on the percent of carbon in leaf tissue perhaps due to stomatal closure and a decrease in photosynthetic activity leading to less assimilation of CO_2_ from the atmosphere (Chen et al. 2015; Ruehr et al. 2009). Surprisingly, the whole community inoculant had a positive interactive effect in drought, perhaps the effects of leaf microbes on C loss during leaf decomposition was dampened (Su, Kuehn, and Phipps 2015). For leaf percent nitrogen and the amount of N relative to C, *B. amyloliquefaciens* had a strong positive effect while the whole community had a negative effect. This enrichment of leaf %N with *B. amyloliquefaciens* indicates that the plant has more resources to contribute to growth and the decrease of %N in the whole community treatment groups may indicate poor nutrient uptake or competition for N with the leaf microbes (Minden and Kleyer 2011; Zhang et al. 2020; Chen et al. 2015). As a part of the root community, *B. amyloliquefaciens* can have a positive effect on the uptake of nitrogen by a plant by volatilization of soil ammonia (Xue et al. 2021) and possibly by N fixation (Ben Abdallah, Frikha-Gargouri, and Tounsi 2018; Cui et al. 2022). The strain used in this study was initially cultured as an endophyte in a previous characterization of the sagebrush phyllosphere (Heil et al., 2024). However, it is possible that the increased nitrogen content of the leaves may be a result of *B. amyloliquefaciens* also inoculating the root microbiome from our spray inoculations, allowing for increased nitrogen availability from the soil.

### Conclusion

The interaction of the leaf microbiome with its plant host is complex and context-dependent, confounding easy explanations and predictions of the effects microbes can have on their host. The clear treatment effects we observed suggest that colonization by non-target microbes do not overshadow their influence on sagebrush seedling physiology and growth. These effects were mixed in their benefit to the plant and often highly dependent on the environmental conditions. We saw a general decrease in photosynthetic activity during drought and with microbial inoculation, however, the size of this effect was less when drought was combined with microbial inoculants. We saw a potential positive effect of inoculation with the bacterium *B. amyloliquefaciens*, with increased leaf nitrogen levels and the highest survival under drought conditions. However, this inoculation also led to a negative effect on total seedling dry weight. Increased water and humidity can contribute to the negative effects of leaf-borne fungal pathogens, and in our experiment this seemed to be augmented by the addition of any microbes, highlighting how microbial interactions can impact effects on a host. As sagebrush is a drought-adapted plant, it is possible that the plant relies on drought as a buffering condition to the negative effects of pathogenic fungi.

## Acknowledgements

This research was supported by a seed grant from the NSF Idaho EPSCoR Program and by the National Science Foundation under the GEM3 award number OIA-1757324. Next-Generation Sequencing was performed by the Argonne National Laboratory. Special thanks to the members of the Bittleston Lab for assistance with data gathering and lab work and to Jennifer Forbey, Marcelo Serpe, and Posy Busby for their guidance and helpful suggestions.

## Competing Interests

None declared.

## Author Contributions

JAH and LSB conceptualized the project. JAH designed the project with essential contributions from LSB, MG, MS, CD, and AA. JAH, MG, MS, CD, AA, and BF acquired the data. JAH analyzed the data. JAH, LSB, and BF contributed to interpretation of the data. JAH drafted the work with contributions from MG, MS, CD, AA, and BF. All authors contributed to revision of the manuscript and approve the final manuscript.

## Data availability

All data are available freely at 10.5281/zenodo.14767517.

## Notes

### Competing Interest Statement

The authors have declared no competing interest.

