## Supplementalinformation for "Context-dependent effects of microbial inoculation on sagebrush seedlings during drought stress"

***Supplemental information for: Context-dependent effects of microbial inoculation on sagebrush seedlings during drought stress***

Table of Contents

Supplemental Methods (page 1-2)

Figure S1 (page 2)

Table S1 (page 3)

Table S2 (page 3)

Table S3 (page 3)

Table S4 (page 4)

Table S5 (page 4)

Table S6 (page 4)

Table S7 (page 5)

Table S8 (page 5)

References (page 5)

Supplemental Methods

*Epiphyte Culturing*

Leaf samples were harvested into sterile microcentrifuge tubes, into which 200 μL of sterilized distilled water was added. The samples were washed by vortexing for 2 minutes, then the entire leaf wash was pipetted onto Potato Dextrose Agar (PDA) plates. Approximately five sterile 4 mm glass beads were added to each plate and gently swirled to distribute the leaf wash across the media surface evenly. The beads were then discarded into a bead waste container. The plates were sealed with parafilm and incubated at room temperature for approximately seven days.

*Endophyte Culturing*

Leaf material was cut into strips (1 mm x 2 mm) using a sterile razor blade and placed into a tea strainer. The samples were surface-sterilized using the set-up shown in Figure 1. The sterilization process involved immersing the samples in a series of baths: 8.3% Tween for 2 minutes, 70% ethanol for 2 minutes, followed by two rinses in sterile water for 2 minutes each (Durán et al., 2021). Tween was used to break surface tension and remove microbial material from the leaf surfaces, ethanol was used to remove Tween and kill surface microbes (epiphytes), and sterile water to wash off residual Tween and ethanol, thereby preventing inhibition of internal microbial growth (endophytes). Finally, using sterilized forceps, the surface-sterilized leaf strips were arranged in a square formation on PDA plates, ensuring adequate spacing between them. The plates were sealed with parafilm and incubated at room temperature for seven days.

*Canopy area measurement*

To measure the leaf canopy area of the sagebrush seedlings we used the Fiji image processing

package version 2.14.0 (Schindelin et al. 2012). Each seedling being inoculated was

photographed in front of a white background with a tape measure extended into frame to provide

scale. Using Fiji, we opened the images and used the Freehand selections tool to trace the bottom

of the plant and make a shape that can be filled with white. We used this technique to separate

the seedling from the pot in the image and make the plant into an isolated object

surrounded by white. We then used the straight-line tool to mark 2 cm on the measuring tape, and

set the scale for the image using the ‘Analyze > Set Scale’ options, entering 2 as the Known

distance. We then changed the image type to 8-bit and binary. After this, we used the wand

selection tool to select the plant, holding down the Shift button while choosing the plant-object

allowed us to also select any smaller isolated plant image fragments that had been separated from

the larger object. We then navigated to the ‘Analyze > Set Measurements’ options and checked the

boxes for Area and Area Fraction. The ‘Analyze > Measure’ options then gave us the area of the

plant-object in the image and the fraction of that area that was not background. We then

multiplied the Area by the Area Fraction and that gave us our canopy measurement.

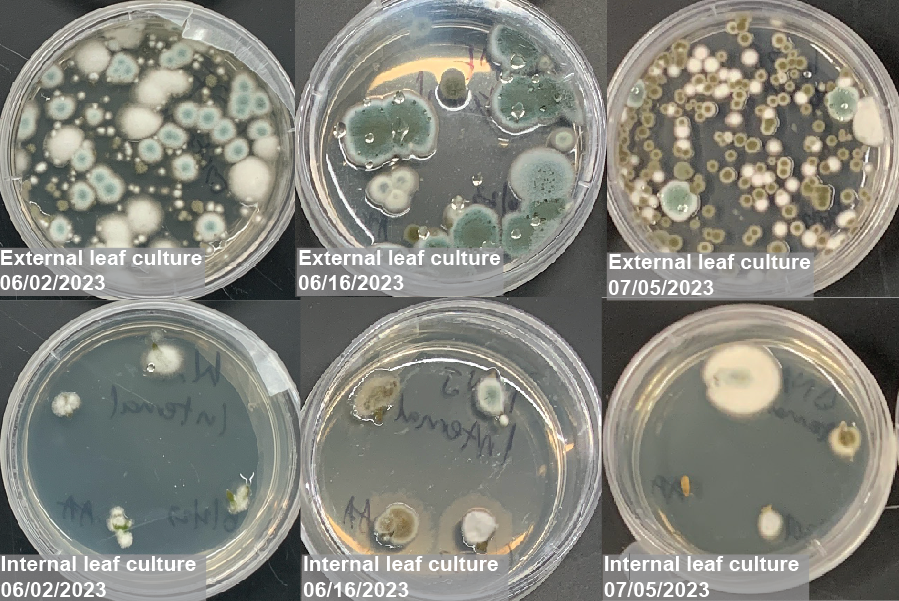

**Figure S1. Example cultures from different time points throughout experiment.** Three fungal morphospecies were found on plants from all test groups throughout the experiment.

**Table S1. Net photosynthesis model results.** We modeled net photosynthesis as the response variable with the explanatory variables watering regime and inoculation including an interactive term and plant individual and date as random intercepts.

```
Family: gaussian
  Links: mu = identity; sigma = identity
Formula: Photo ~ Watering * inoculant + (1 | Plant_ID) + (1 | Date)

Multilevel Hyperparameters:
~Date (Number of levels: 6)
              Estimate Est.Error l-95% CI u-95% CI Rhat Bulk_ESS Tail_ESS
sd(Intercept)     3.68      1.66     1.66     7.88 1.00     2487     3223

~Plant_ID (Number of levels: 80)
              Estimate Est.Error l-95% CI u-95% CI Rhat Bulk_ESS Tail_ESS
sd(Intercept)     0.70      0.50     0.03     1.85 1.00     3020     3047

Regression Coefficients:
                        Estimate Est.Error l-95% CI u-95% CI  Rhat
Intercept                 10.98      2.32     6.29    15.43   1.00
Wateringdrought           -0.81      2.33    -5.35     3.71   1.00
inoculantsterilewater      7.67      2.00     3.81    11.56   1.00
inoculantb.amy            -0.07      2.01    -3.96     3.85   1.00
inoculantwholecommunity   -3.81      1.99    -7.66     0.05   1.00
drought:sterilewater      -5.23      2.74   -10.61     0.13   1.00
drought:b.amy             -0.43      2.74    -5.78     4.89   1.00
drought:wholecommunity     2.79      2.73    -2.50     8.14   1.00
```

**Table S2. Canopy area model results.** We modeled canopy area as the response variable with the explanatory variables watering regime and inoculation including an interactive term.

```
Family: gaussian
  Links: mu = identity; sigma = identity
Formula: CA ~ Watering * inoculant

Regression Coefficients:
                          Estimate Est.Error l-95% CI u-95% CI Rhat
Intercept                  8.40      1.41     5.63    11.25    1.00
Wateringdrought           -0.80      2.01    -4.84     3.15    1.00
inoculantsterilewater      3.12      2.01    -0.81     7.02    1.00
inoculantb.amy             0.40      2.18    -3.83     4.65    1.00
inoculantwholecommunity   -4.16      2.01    -8.19    -0.11    1.00
drought:sterilewater      -1.46      2.84    -7.06     4.02    1.00
drought:b.amy             -0.75      2.98    -6.52     5.20    1.00
drought:wholecommunity     1.76      2.80    -3.95     7.21    1.00
```

**Table S3. Whole plant dry weight model results.** We modeled whole plant dry weight as the response variable with the explanatory variables watering regime and inoculation including an interactive term.

```
Family: gaussian
  Links: mu = identity; sigma = identity
Formula: Dry_Weight ~ Watering * inoculant

Regression Coefficients:
                          Estimate Est.Error l-95% CI u-95% CI Rhat
Intercept                  0.40      0.07     0.25     0.54    1.00
Wateringdrought            0.24      0.10     0.04     0.44    1.00
inoculantsterilewater     -0.05      0.10    -0.25     0.14    1.00
inoculantb.amy            -0.13      0.10    -0.33     0.07    1.00
inoculantwholecommunity   -0.23      0.10    -0.43    -0.04    1.00
drought:sterilewater      -0.07      0.14    -0.36     0.21    1.00
drought:b.amy             -0.10      0.14    -0.38     0.18    1.00
drought:wholecommunity     0.16      0.14    -0.12     0.44    1.00
```

**Table S4. Root length model results.** We modeled root length as the response variable with the explanatory variables watering regime and inoculation including an interactive term.

```
Family: gaussian
  Links: mu = identity; sigma = identity
Formula: Root_Length_cm ~ Watering * inoculant

Regression Coefficients:
                          Estimate Est.Error l-95% CI u-95% CI Rhat
Intercept                 17.58      1.12    15.37    19.72    1.00
Wateringdrought           -0.79      1.57    -3.81     2.20    1.00
inoculantsterilewater     -1.35      1.56    -4.33     1.82    1.00
inoculantb.amy            -2.34      1.58    -5.31     0.81    1.00
inoculantwholecommunity   -5.23      1.63    -8.39    -2.00    1.00
drought:sterilewater       2.21      2.22    -2.19     6.47    1.00
drought:b.amy             -1.60      2.20    -6.01     2.68    1.00
drought:wholecommunity     4.31      2.28    -0.15     8.78    1.00
```

**Table S5. Stem length model results.** We modeled stem length as the response variable with the explanatory variables watering regime and inoculation including an interactive term.

```
Family: gaussian
  Links: mu = identity; sigma = identity
Formula: Stem_Length_cm ~ Watering * inoculant

Regression Coefficients:
                           Estimate Est.Error l-95% CI u-95% CI Rhat
Intercept                  5.56      0.41     4.78     6.38     1.00
Wateringdrought           -0.57      0.59    -1.68     0.57     1.00
inoculantsterilewater      0.74      0.58    -0.39     1.92     1.00
inoculantb.amy             0.11      0.58    -1.02     1.27     1.00
inoculantwholecommunity   -1.01      0.60    -2.21     0.15     1.00
drought:sterilewater       0.44      0.83    -1.19     2.08     1.00
drought:b.amy              0.04      0.83    -1.64     1.65     1.00
drought:wholecommunity     0.29      0.84    -1.34     1.93     1.00
```

**Table S6. Plant weight %C model results.** We modeled plant weight %C as the response variable with the explanatory variables watering regime and inoculation including an interactive term.

```
Family: beta
  Links: mu = logit; phi = identity
Formula: Weight_percC * 0.01 ~ Watering * inoculant

Regression Coefficients:
                          Estimate Est.Error l-95% CI u-95% CI Rhat
Intercept                 -0.35      0.03    -0.41    -0.28    1.00
Wateringdrought           -0.09      0.04    -0.18    -0.00    1.00
inoculantsterilewater      0.02      0.04    -0.07     0.11    1.00
inoculantb.amy             0.00      0.04    -0.08     0.09    1.00
inoculantwholecommunity   -0.03      0.04    -0.12     0.05    1.00
drought:sterilewater       0.03      0.06    -0.09     0.16    1.00
drought:b.amy              0.06      0.06    -0.06     0.19    1.00
drought:wholecommunity     0.15      0.06     0.03     0.27    1.00
```

**Table S7. Plant weight %N model results.** We modeled plant weight %N as the response variable with the explanatory variables watering regime and inoculation including an interactive term.

```
Family: beta
  Links: mu = logit; phi = identity
Formula: Weight_percN * 0.01 ~ Watering * inoculant

Regression Coefficients:
                          Estimate Est.Error l-95% CI u-95% CI Rhat
Intercept                 -3.29      0.05    -3.38    -3.20    1.00
Wateringdrought            0.01      0.06    -0.12     0.14    1.00
inoculantsterilewater     -0.03      0.07    -0.16     0.09    1.00
inoculantb.amy             0.15      0.06     0.02     0.27    1.00
inoculantwholecommunity   -0.09      0.07    -0.21     0.05    1.00
drought:sterilewater       0.03      0.09    -0.15     0.21    1.00
drought:b.amy             -0.09      0.09    -0.27     0.09    1.00
drought:wholecommunity     0.07      0.09    -0.11     0.25    1.00
```

**Table S8. C:N model results.** We modeled C:N as the response variable with the explanatory variables watering regime and inoculation including an interactive term.

```
Family: gamma
  Links: mu = log; shape = identity
Formula: CNratio ~ Watering * inoculant

Regression Coefficients:
                          Estimate Est.Error l-95% CI u-95% CI Rhat
Intercept                  2.46      0.04     2.39     2.54    1.00
Wateringdrought           -0.07      0.05    -0.18     0.04    1.00
inoculantsterilewater      0.04      0.05    -0.07     0.14    1.00
inoculantb.amy            -0.14      0.05    -0.25    -0.04    1.00
inoculantwholecommunity    0.07      0.05    -0.04     0.17    1.00
drought:sterilewater       0.00      0.08    -0.15     0.15    1.00
drought:b.amy              0.13      0.08    -0.02     0.28    1.00
drought:wholecommunity     0.02      0.08    -0.13     0.17    1.00
```
